# Spinal Cord Stimulation using time-dynamic pulses achieves faster and longer reversal of allodynia compared to tonic pulses in a rat model of neuropathic pain

**DOI:** 10.1101/2023.03.07.531522

**Authors:** Muhammad Edhi, Changfang Zhu, Ki-Soo Jeong, Victoria Rogness, Rosana Esteller, Carl Saab

## Abstract

Spinal cord stimulation (SCS) utilizing time-dynamic pulses (TDPs) is an emergent field of neuromodulation that continuously and automatically modulates pulse parameters. We previously demonstrated that TDPs delivered for 60 min at sub-paresthesia amplitudes significantly reversed allodynia in a rat model of neuropathic pain. Because we observed these anti-allodynic effects persisted post-cessation, we investigated the extended temporal dynamics of SCS-induced analgesia. We hypothesized that TDPs achieve a longer duration of analgesia than tonic stimulation. Both TDPs and tonic stimulation reversed PWT to near pre-chronificiation levels within 30 minutes. Most TDPs exhibited significantly slower ramp-up slope (analgesia ‘wash-in’ rates) compared to tonic stimulation (amplitude modulation: 0.16±0.03 min^-1^, pulse width modulation: 0.18±0.05 min^-1^, stochastic modulation: 0.17±0.04 min^-1^, tonic: 0.31±0.06 min^-1^). All TDPs showed slower wind-down slopes (analgesia ‘wash-out’ rates) compared to tonic (-0.29±0.07 min^-1^), with pulse width modulation (-0.11±0.02 min^-1^) reaching significance. Extending SCS from 60 to 90 minutes revealed all TDPs maintain analgesic efficacy longer than tonic stimulation, which decreased significantly at both 75 and 90 minutes (from 13.8±0.5 g to 12.3±0.9 g and to 11.0±0.5 g, respectively). Although TDPs and tonic stimulation comparably mitigated allodynia, TDPs generally exhibited slower temporal dynamics, suggesting longer-lasting analgesic effects and potentially different mechanisms of action.

## Introduction

Neuromodulation is the manipulation of neural activity through targeted delivery of electromagnetic stimuli or pharmacological agents that modulate the activity of the nervous system, thus offering a wide range of modalities for the treatment for neurological diseases, including chronic pain. Electrical spinal cord stimulation (SCS) refers to an FDA-approved treatment for chronic and neuropathic pain conditions that involves delivering electric current via epidural leads implanted into the spinal cord. This method of treatment is thought to attenuate pain perception via blocking the transmission of nociceptive signaling in the spinal cord dorsal horn according to the Gate Control Theory.[1,2] Conventional SCS, which delivers currents via static electrical pulses with constant stimulation parameters (i.e. ‘tonic’), may introduce paresthesias such as tingling sensations at certain stimulus intensities above perception threshold.[3] More recent SCS paradigms that deliver electrical stimuli below perceptual levels (sub-paresthesia) have also been shown to achieve analgesic efficacy. Moreover, SCS using time-dynamic pulses (TDPs) is an emerging field of neuromodulation and a promising advancement that, unlike tonic stimulation, modulates stimulation pulse parameters with signals that change continuously and automatically.[4–6]

Our team previously demonstrated that TDPs delivered at sub-paresthesia amplitudes significantly reverse allodynia in a rat model of neuropathic pain.[6] Specifically, we showed that different TDP stimulation consisting of pulse width modulation, amplitude modulation, sinusoidal rate modulation (abbreviated here as rate), and stochastic modulation delivered for 60 min at sub-paresthesia amplitudes significantly reversed allodynia in rats two weeks after chronic constriction injury (CCI). Moreover, SCS reversed an electroencephalography (EEG) signature of spontaneous pain in rats with CCI.[6,7] Our results also indicated that TDPs demonstrated sustained attenuation of hypersensitivity at the end of the 60 min SCS period, suggesting that the analgesic effects of TDPs may extend beyond the hour-long duration tested. In this study, therefore, we evaluated the extended temporal dynamics of the analgesic properties of SCS. We hypothesized that TDPs achieve a longer duration of analgesia (i.e. therapy wash-out) compared to tonic stimulation. Therefore, we sought to compare the efficacy and time course of analgesic responses to TDP modulation in amplitude, pulse width, and rate to that of tonic stimulation in the same animal model of CCI. Compared to our previous study, here we extended the SCS period to 90 min for a more comprehensive temporal assessment of the time producing (i.e., during SCS ‘on’) and maintaining analgesic efficacy (i.e., after SCS ‘off’), including the time to onset and decay of PWT that mirror the wash-in time to induce analgesia and the wash-out time to lose analgesia. Our results generally support our hypothesis that TDPs achieve longer durations of analgesia during SCS ‘on’ and after SCS ‘off’ compared to tonic stimulation.

## Methods

We conducted a longitudinal, randomized, double-blind crossover experimental design in n=23 rats, including eight previously reported samples[6], and additional fifteen new samples (Fig 1). For the new samples (n=15), we extended the SCS duration from 60 to 90 minutes. Behavioral responses to von Frey filaments during SCS were recorded as paw withdraw threshold (PWT) at 15 min intervals. Each rat received five sessions of SCS assigned in a random order: 4 different TDP stimulation sessions and one tonic stimulation session. The response curves of PWT per rat and per SCS session were fitted with a double sigmoidal function (with average fitting deviation < 15%, calculated as a ratio of summed square deviation vs summed square target data, Fig 2). We then characterized the time course of PWT responses for the time to reach half maximum PWT reversal during SCS ‘on’ and the decay time to half maximum PWT reversal after SCS ‘off’.

**Figure 1.**
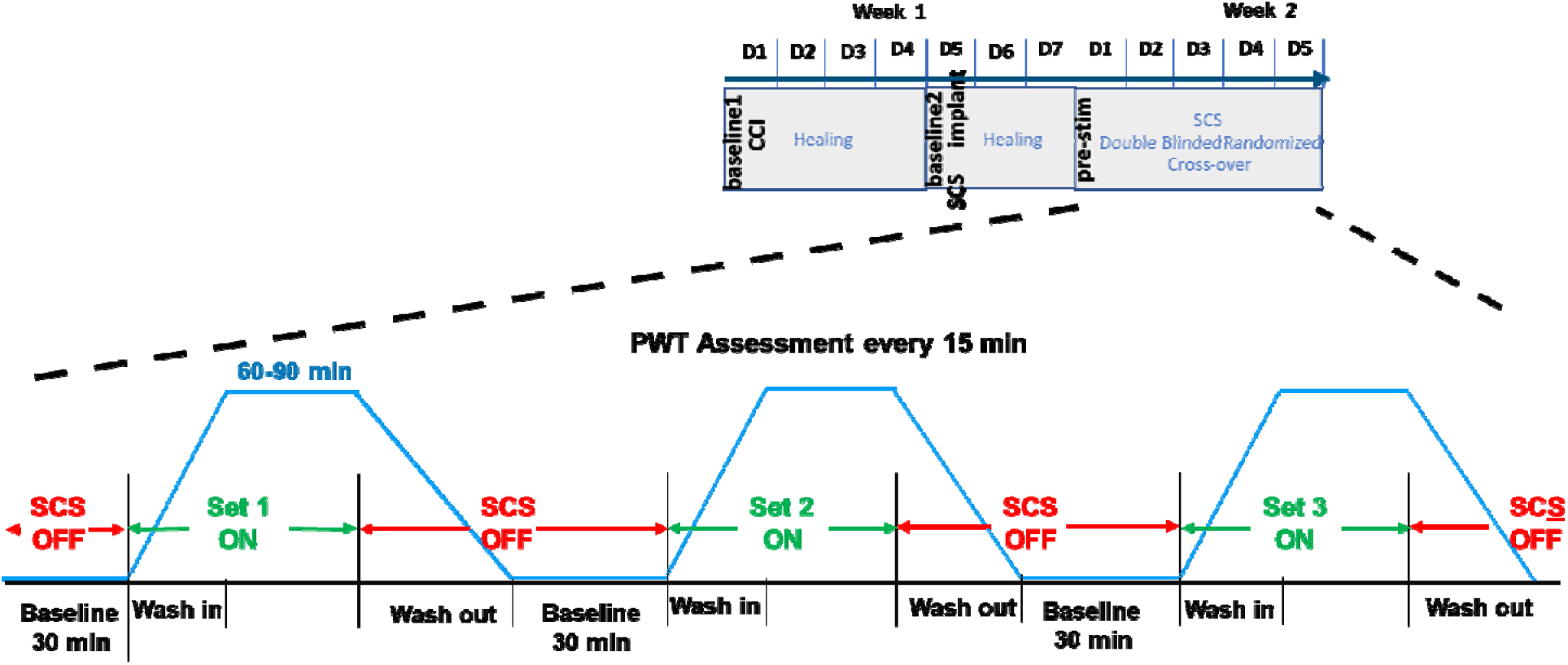
Time course of behavioral paw withdrawal threshold (PWT) measurements in relation to chronic constriction injury (CCI) and spinal cord stimulation (SCS).

**Figure 2.**
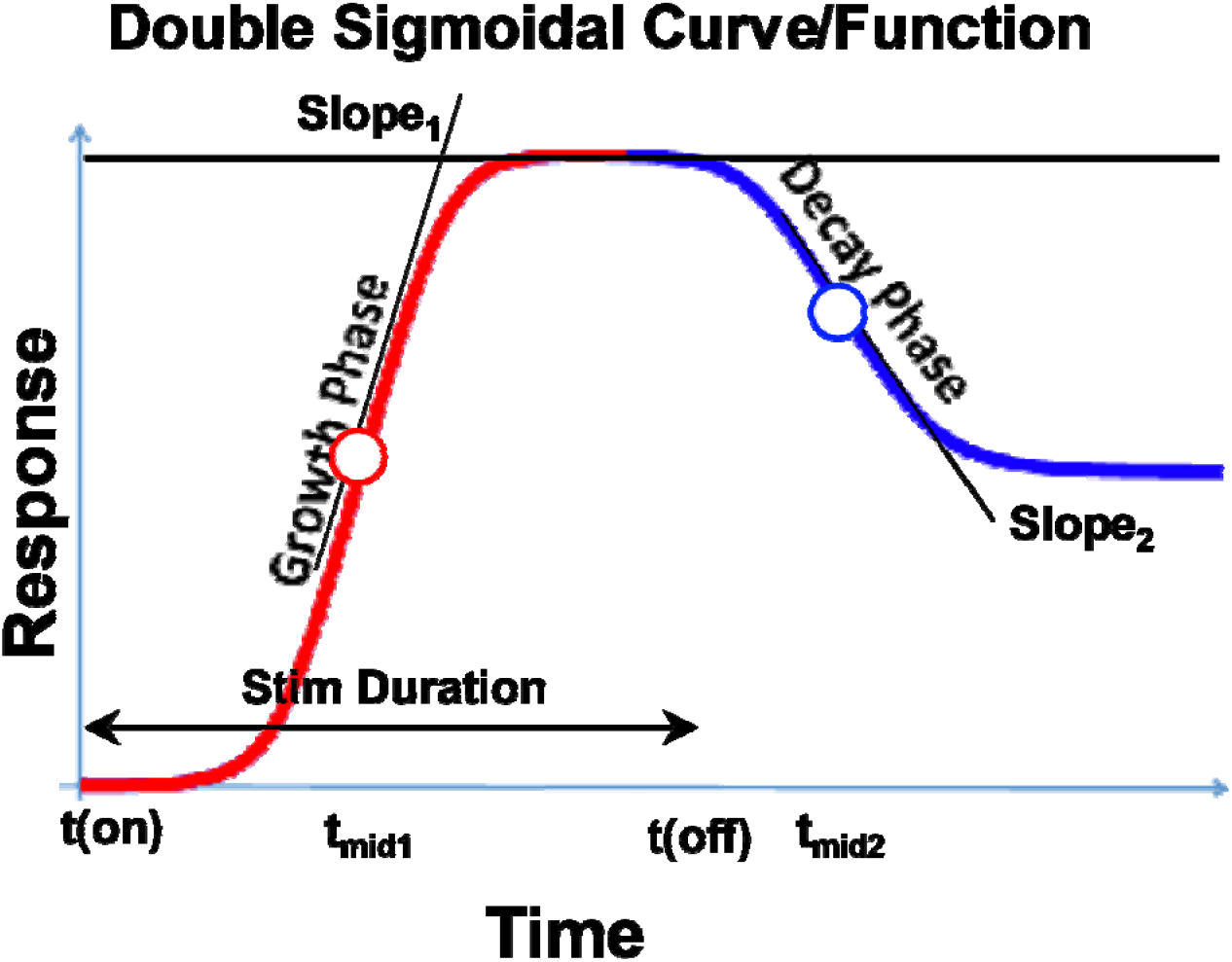
Double sigmoidal curve function for modeling response. t_mid1_, t_mid2_, full-width half-maxima (FWHM), slope_1_, and slope_2_ all indicate different metrics for gauging response onset and decay. t_mid1_ and t_mid2_ indicate time for response to rise and to decay to half maxima since the start and end of stimulation, respectively. FWHM reflects the percent of stimulation time with response above half maximum. Slope_1_ and slope_2_ determine the speed of growth and speed of decay, respectively.

### Experimental animals and surgical operations

Adult (200–300 g) male Sprague Dawley rats were housed under a 12-h light/dark cycle in a temperature and humidity-controlled environment. Food and water were available ad libitum. All surgical procedures were performed under deep anesthesia (isoflurane, 2–2.5%). All the methods were carried out in accordance with the relevant guidelines and regulations. Experiments were approved by Cleveland Clinic Lerner Research Institute Institutional Animal Care and Use Committee (IACUC) and, for the first cohort, by Rhode Island Hospital IACUC. Induction of CCI, implantation of the spinal leads, and configuration and delivery of SCS all followed methods and procedures outlined in our previous study.[6]

### Responsive curve analysis and temporal evaluation of PWT time course

Paw withdrawal threshold to von Frey filaments were recorded during SCS ‘on’ (60 or 90 min) and ‘off’ (30 minutes) at 15 min intervals. The time course of PWT reflected the analgesic effect of SCS, which we assumed follows a biphasic ramp-up after SCS onset and wind-down after SSC ‘off’. Hence, we identified the baseline, maximum, and residual PWT responses, and the corresponding timeline when these responses occurred. We adapted a double-sigmoidal model to fit the PWT responses over time using optimization functions provided in MATLAB (Mathworks, MA). Sigmoidal and double sigmoidal curves have been widely used to model dose-response data or biological growth data.[8,9] In our study, the five different types of SCS patterns represent different treatments and the two variable durations of SCS (60 vs. 90 min) represent the dose, while the PWT reversal represents response to treatment.

The target function of the sigmoidal curve fitting was the time course of PWT change during SCS ‘on’ and after SCS ‘off.’ To reduce the difference in scale and range of PWT values due to variations in baseline and maximum PWT reversal, we pre-processed PWT(t) as following: PWT values were first normalized to baseline prior to SCS ‘on’ (t = 0), and then had baseline subtracted such that the response prior to SCS onset was set to zero. This normalization procedure removed variation in baseline across individual animals, and the subtraction yielded absolute changes from baseline. Most response curves were scaled roughly to a range within [0, 2] post pre-processing. The pre-processed PWT data served as the target function *f*_target_(t)

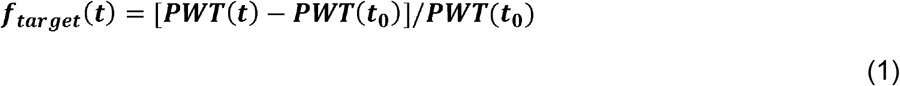

The fitting curve was represented as a time dependent function *f*_fit_ (t):

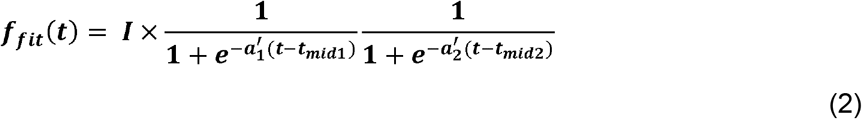

where (I) was a scaling factor to account for the magnitude of PWT reversal (response); t_mid1_ estimates the time at which the response has risen to half of its maximum; t_mid2_ estimates the time at which the response has decayed halfway from its maximum to its final value; and the parameters a_1_ and a_2_ determine the steepness of the ramp-up and wind-down phases of the sigmoidal curves, respectively, but do not exactly equal to the slopes of the curve at times t_mid1_ and t_mid2_. The latter was represented as slope_1_ and slope_2_ and calculated from the 1^st^ order derivative of the double-sigmoidal function *f*_fit_ (t) at times t_mid1_ and t_mid2_. These metrics represent the ramp-up and wind-down effects, respectively, of analgesia as measured by PWT.

The fitting parameters were obtained through minimizing the cost function, which was defined as the summed squared deviation (SSD) of the fit data points to the raw data points via:

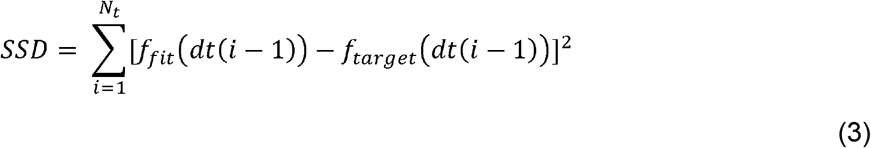

where *dt* refers to the time interval between observations (*dt* = 15 min), and *N_t_* refers to the total number of observations during recording period (*N_t_* = 7 for the first cohort of 8 samples and *N_t_* = 9 for the second cohort of 15 samples).

The quality of fitting was evaluated by calculating the percent deviation which is the ratio of the summed square of deviation over the summed square of raw data,

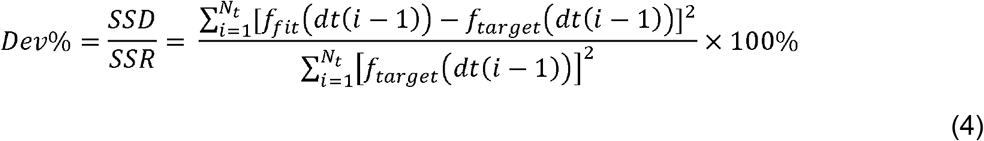

where the summed square of the raw data was calculated as:

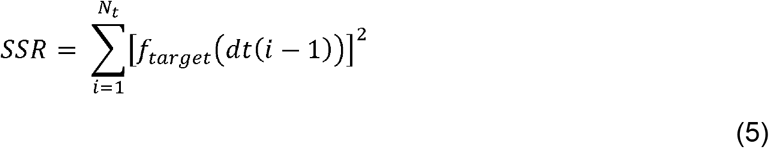

From these fitted models, various temporal metrics were calculated, including the fullwidth-half-maximum (FWHM) of the response, which was defined as the percentage of time with response above half of the maximal PWT recovery, and calculated as the duration between t_mid1_ and t_mid2_, normalized by the total duration of SCS stimulation, i.e. (t_mid2_ – t_mid1_)/T_stim_, where T_stim_ = 60 min for the first cohort of 8 samples and T_stim_ = 90 min for the second cohort of 15 samples (Table 1).

**Table 1.**
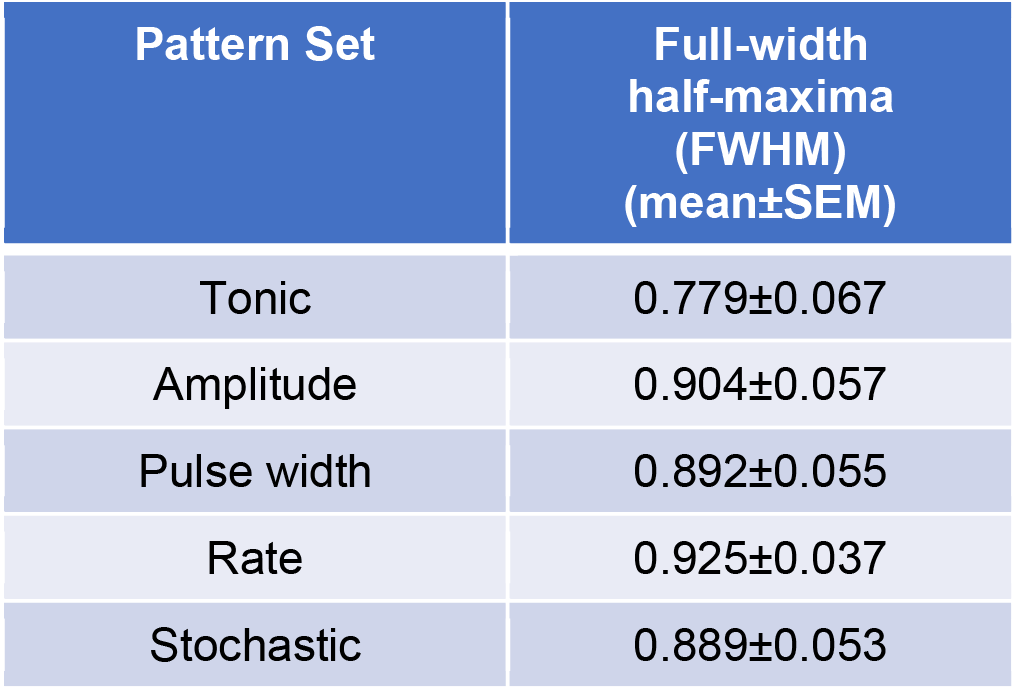
Full-width half-maxima (FWHM) by pattern set across both cohorts (N=23), which reflects the percentage of the total time analgesic response is demonstrated for a given pattern. FWHM for TDP stimulation were closer relative to each other than to tonic stimulation (Mean±SEM).

| Pattern Set | Full-width half-maxima (FWHM) (mean±SEM) |
| --- | --- |
| Tonic | 0.779±0.067 |
| Amplitude | 0.904±0.057 |
| Pulse width | 0.892±0.055 |
| Rate | 0.925±0.037 |
| Stochastic | 0.889±0.053 |

### Statistical analysis

We performed our statistical analysis with native functions in Prism 9 (GraphPad Software, San Diego). We considered α= 0.05 as statistically significant (*). Additionally, as we sought to compare TDPs collectively against tonic stimulation as the gold standard (rather than between-pattern comparison, for example amplitude vs. pulse width TDP), we implemented a two-way repeated measures omnibus test (ANOVA for paw withdrawal threshold, mixed-effects model for the double-sigmoidal slope values), followed by multiple comparisons without post hoc corrections for behavioral analysis of PWT data. For each pulse pattern, we assessed the time course of PWT under the stimulation using the same pattern across all rats by aggregating the data for which both cohorts received stimulation for a same period after onset (post-SCS, 0 < t ≤ 60) and then performed statistical tests to compare the PWT at each time point post stimulation onset to that obtained at the time point pre-SCS (t=0).

### Data availability

The datasets generated and/or analyzed during the current study are available from the corresponding authors on reasonable request.

## Results

### Sigmoidal curve fitting

We excluded nine out of 115 datasets for which either: 1) the percent deviation exceeded 50% (n = 2); or 2) the scaling factor (I) was smaller than 0.1 (in which the fitted model demonstrates a change in PWT response of less than 10% from baseline, n = 1), or negative (in which the fitted model ran in the opposite direction of the expected change in PWT response, n = 3); or 3) both aforementioned exclusion criteria were met (n = 3). No rat had greater than two data points excluded. The average percent deviation of fitting for all models (including all samples without exclusions) fell under 15% (tonic: 7.9±2.1%; amplitude: 15.2±4.3%; pulse width: 13.4±4.2%; rate: 13.6±4.2%; stochastic: 12.1±3.4%) (see Supplementary Fig. 2), and the average percent deviation of fitting after exclusion fell under 10% (except 10.5% for amplitude).

### SCS modulation of PWT

All five SCS patterns reduced allodynia towards pre-CCI values, such that PWT were significantly elevated at t=30, 45 and 60 min compared to t=0 (Fig 3), according to the following timeline: For tonic stimulation, PWT significantly increased from 5.9±0.3 g at t=0 to 7.2±0.5 g at t=15 min (p<0.05), 11.2±0.6 g at t=30 min (p<0.001), 10.9±0.8 g at t=45 min (p<0.001) and 11.7±0.8 g at t=60 min (p<0.001). For TDP amplitude modulation, PWT significantly increases from 6.0±0.3 g at t=0 to 7.5±0.4 g at t=15 min (p<0.01), 9.7±0.5 g at t=30 min (p<0.001), 9.1±0.6 g at t=45 min (p<0.001) and 9.6±0.6 g at t=60 min (p<0.001). For TDP pulse width modulation, PWT significantly increased from 6.3±0.5 g at t=0 to 9.4±0.7 g at t=30 min (p<0.01), 9.1±0.7 g at t=45 min (p<0.001), and 10.7±0.8 g at t=60 min (p<0.001). However, the increase in PWT during pulse width modulation did not reach significant levels at t=15 min compared to t=0. For TDP rate modulation, PWT significantly increased from 5.8±0.3 g at t=0 to 8.0±0.4 g at t=15 min (p<0.001), 10.4±0.5 g at t=30 min (p<0.001), 11.2±0.6 g at t=45 min (p<0.001) and 10.2±0.5 g at t=60 min (p<0.001). For TDP stochastic modulation, PWT significantly increased from 6.3±0.5 g at t=0 to 8.2±0.4 g at t=15 min (p<0.01), 9.1±0.4 g at t=30 min (p<0.001), 9.5±0.5 g at t=45 min (p<0.001), and 10.3±0.5 g at t=60 min (p<0.001). For each timepoint between t=0 and 60 min at 15 min intervals, we compared PWT during the 4 TDP stimulation versus tonic stimulation (Supplemental Fig 1), whereby the increase in PWT was not significantly different at t=0 and t=15 min. However, PWT was significantly higher during tonic stimulation compared to TDP stimulation, in particular at t=30 min after pulse width (9.4±0.7g, p<0.05) and stochastic modulation (9.1±0.4 g, p<0.01) compared to tonic stimulation (11.2±0.6 g), at t=45 min after amplitude (9.1±0.6 g, p<0.05) and pulse width modulation (9.1±0.7 g, p<0.05) compared to tonic stimulation (10.9±0.8 g), and at t=60 min after amplitude (9.6±0.6 g, p<0.05) and rate modulation (10.2±0.5 g, p<0.05) compared to tonic stimulation (11.7±0.8 g). No single TDP stimulation achieved consistently higher analgesic efficacy relative to tonic stimulation.

**Figure 3.**
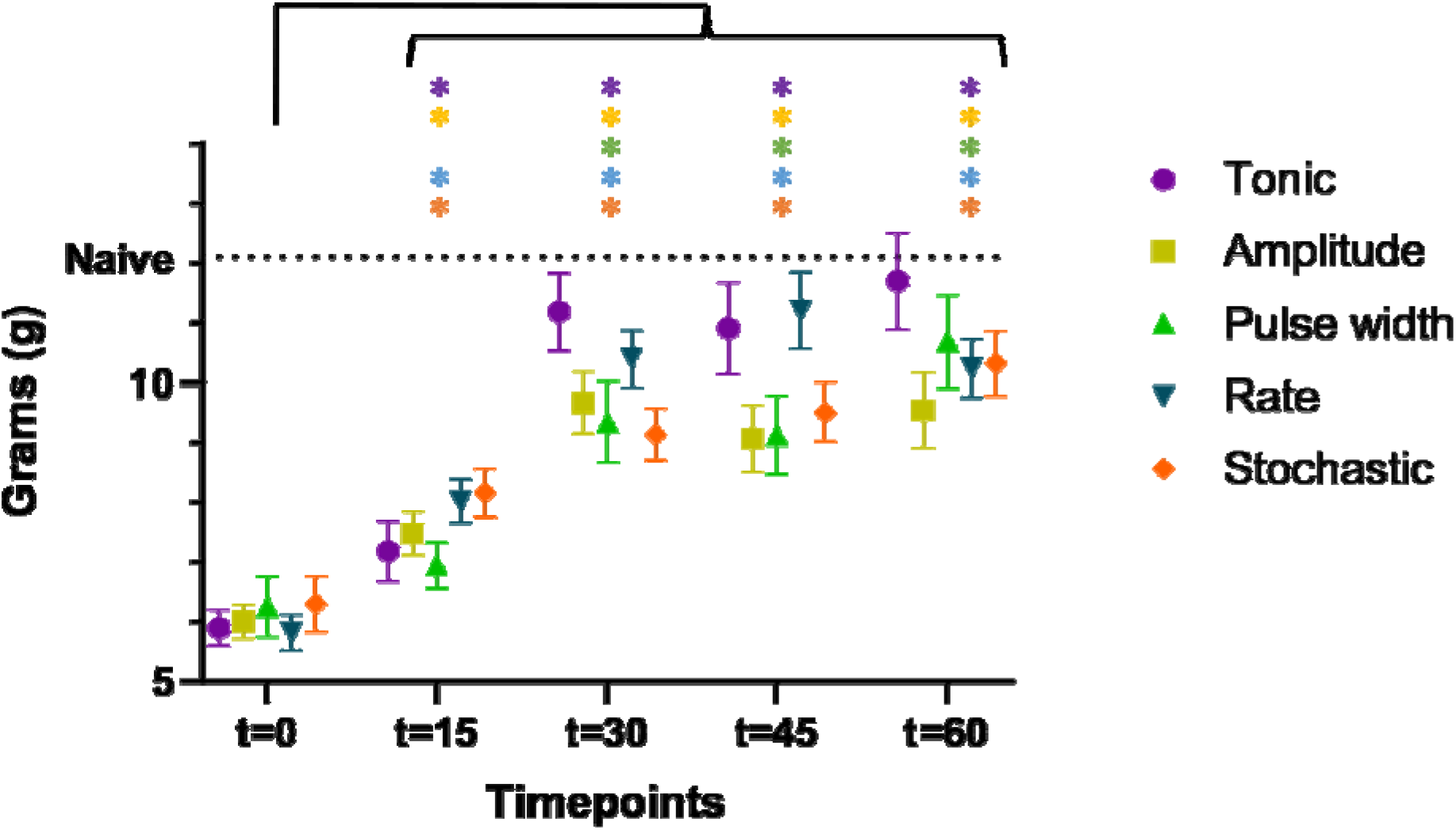
Paw withdrawal threshold (PWT) time course during 5 different patterns of SCS (n=23 rats).

### Impact of extending SCS duration from 60 to 90 min on PWT

All TDP stimulation maintained comparable anti-allodynic effects at t=60, 75 and 90 min (Fig 4). However, these effects significantly degraded for tonic stimulation at t=75 min (12.3±0.9 g, p< 0.05) and further at t=90 min (11.0±0.5 g, p< 0.01) compared to t=60 min (13.8±0.5 g).

**Figure 4.**
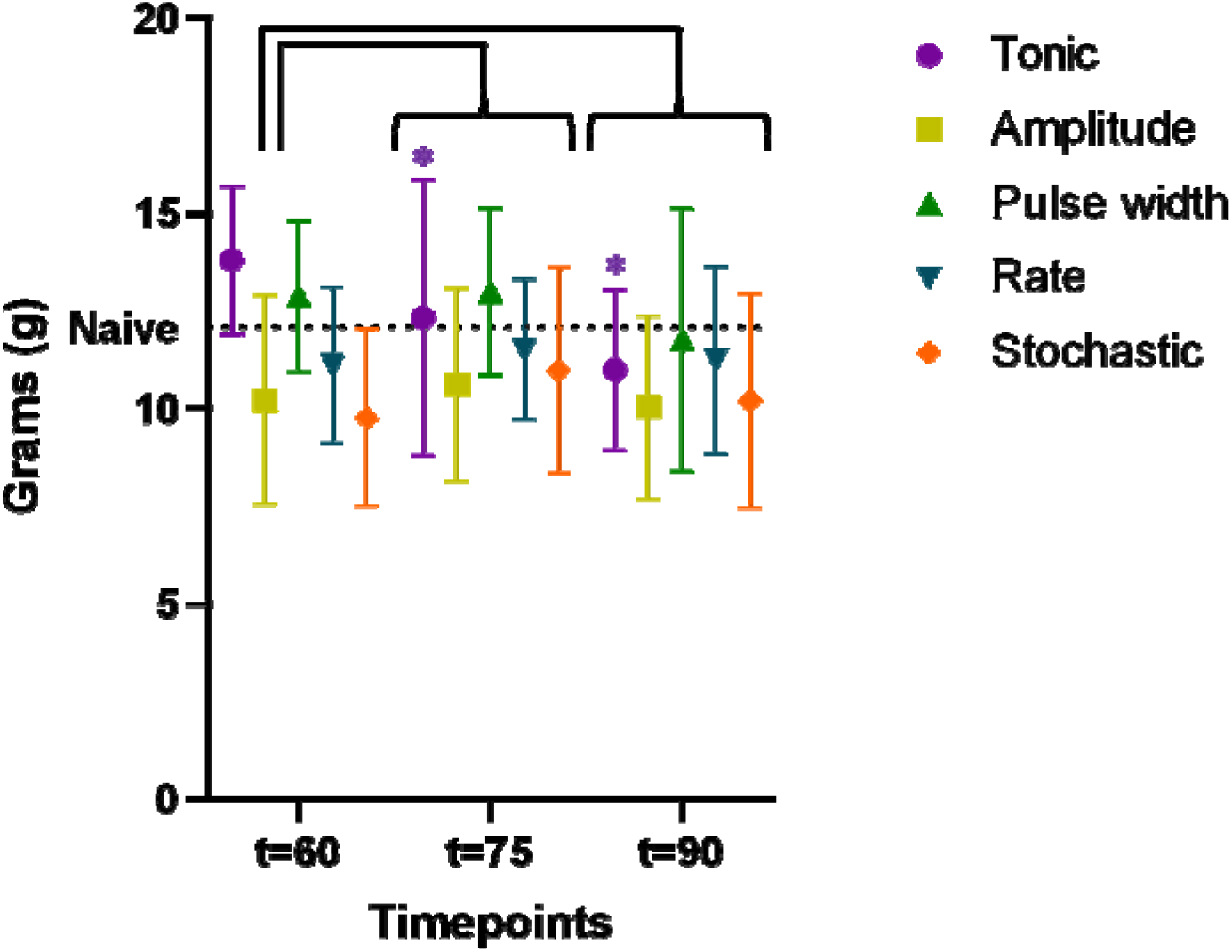
Paw withdrawal threshold (PWT) time course for extended stimulation period up to t=90 min (n=15 rats). Only tonic SCS exhibited significantly degraded efficacy at t=75 min, with further decline at t=90 min, whereas PWT during TDPs remained stable during the same time period.

### Wash-in and wash-out effects on PWT

To evaluate the time course of ramp-up and wind-down of the anti-allodynic effects, we compared slope values for each TDP stimulation to tonic stimulation, whereby a greater (more positive) slope_1_ indicated a faster wash-in and a greater (less negative) slope_2_ indicated a slower wash-out. Slope_1_ values (Fig 5A) were significantly lower for amplitude modulation (0.16±0.03 min^-1^, p<0.05), pulse width modulation (0.18±0.05 min^-1^, p<0.05) and stochastic modulation (0.17±0.04 min^-1^, p<0.05) compared to tonic stimulation (0.31±0.06 min^-1^). However, slope_1_ for rate modulation (0.33±0.09 min^-1^) was not different from that of tonic stimulation. Slope_2_ values (Fig 5B) were higher (i.e., *slower* washout) for amplitude modulation (−0.12±0.02 min^-1^), rate modulation (−0.17±0.04 min^-1^) and stochastic modulation (−0.21±0.06 min^-1^) compared to tonic stimulation (−0.29±0.07 min^-1^), with pulse width modulation reaching significance (−0.11±0.02 min^-1^, p<0.05). Mean FWHM of the response curve for each TDP stimulation was higher compared to tonic stimulation, although the difference did not reach significance (Table 1), noting that FWHM values for TDP stimulation were closer in range relative to tonic stimulation.

**Figure 5.**
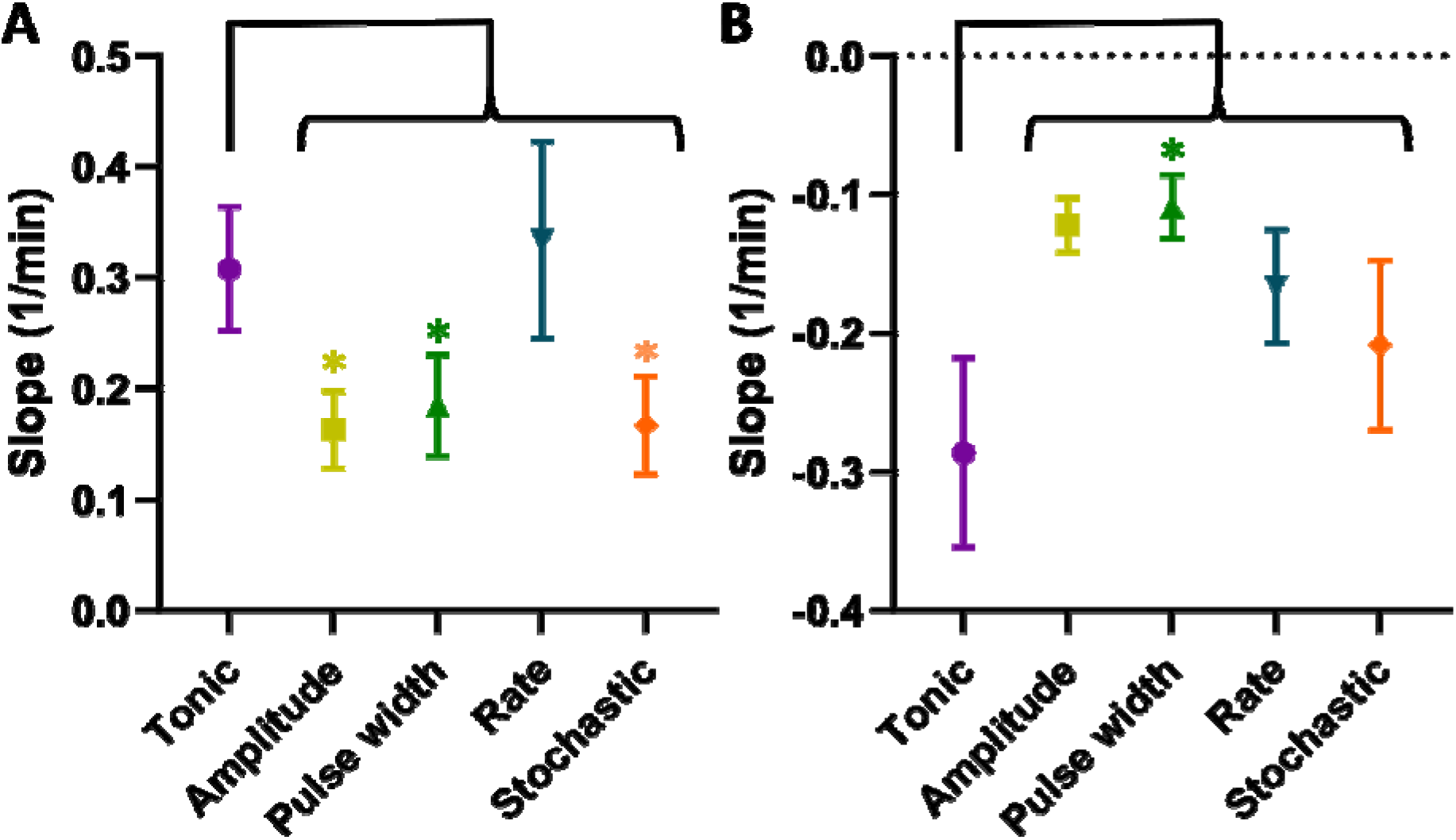
A) Slope1 (speed of growth or “wash-in”). TDPs (amplitude, pulse width, and stochastic rate) had significantly slower wash-in times compared to tonic stimulation, with rate modulation achieving the fastest speed. B) Slope2 (speed of decay or “wash-out”). TDP stimulation had slower wash-out times compared to tonic SCS, with pulse width reaching significance and having the longest wash-out.

## Discussion

In this study, we investigated the anti-allodynic effects of sub-paresthesia SCS consisting of four dynamically modulated pulse patterns (TDPs) compared to conventional tonic stimulation in a rat model of neuropathic pain, using PWT as a surrogate measure of analgesic efficacy. One advantage of this study is the double-blind, cross-over design with a randomized testing order, enabling a more robust comparison of different pulse patterns of stimulation. Consistent with our previous study, our results demonstrate that TDPs and tonic stimulation significantly reversed allodynia within thirty minutes of SCS onset and throughout the duration of SCS lasting 60 min. Overall, analgesic efficacy was higher during tonic stimulation between 30 and 60 min, however it significantly degraded gradually for tonic stimulation at 75 and 90 min compared to all four TDP stimulation, whose analgesic efficacy was maintained over the same extended period. These findings may reflect tolerance to tonic stimulation or desensitization of the neural circuits in the spinal cord to ongoing, constant pulse stimulation[10,11], whereas dynamic neuromodulation, such as the stimulation patterns offered by TDPs, may overcome this phenomenon.

Further investigation into the temporal dynamics of analgesic efficacy (using a novel model fitting method to approximate PWT based on a sigmoidal curve function) revealed delayed, but prolonged analgesic effects for TDPs relative to tonic stimulation. Moreover, FWHM values for all TDP stimulation exceeded those of tonic stimulation and compared favorably with each other, suggesting that TDP stimulation effectively sustain analgesia, while bolstering the idea of distinct mechanisms between TDPs and tonic stimulation. The sigmoidal function has been widely used to describe the dose-response relationship or growth patterns in many physiological or pharmacological studies, as it enables the evaluation of an induced effect in a biological system as a function of the level of the stimulus.[9] The benefits of approximating the discrete assessments of PWT with fitted mathematical models were four-fold: 1) it smoothed the variations in behavioral responses due to the limited temporal resolution and the noise inherent to animal behavior; 2) it allowed quantitative estimates for depicting the time course of said responses; 3) it enabled comparisons between two sets of data from distinct cohorts, which had different numbers of time points due to differences in dosing regimens; and 4) it provided a mathematical tool to identify outliers. Such methods have been applied to assess clinical analgesia and the effects of SCS in animal models.[12–14] The overall percent deviation (< 10%) suggested that the selected model reasonably resembled the individual time courses of PWT. However, we acknowledge the limitation that our model might not represent a mathematically perfect fit, and that other models could be employed in future studies.

Whereas the mechanisms of action for tonic and virtually all forms of SCS remains incompletely understood, the recruitment of mechanical Aβ-fiber afferents likely “gates” the transmission of nociceptive signals.[2,15,16] Given that only tonic stimulation elicited tolerance over prolonged stimulation periods in our study, it is possible that TDP-modulated SCS more closely replicate the irregular and asynchronous activity in physiological states of the dorsal horn.[17–19] For example, amplitude and pulse width-modulated stimulation potentially mimic the physiological encoding of mechanosensory stimuli.[20,21] Meanwhile, rate-modulated stimulation may replicate the firing rate of slowly adaptive primary afferents, thereby altering the reflex response.[18] It may also induce long-term potentiation mediated by NMDA receptors or glial cells.[22–24] Investigation of these and other mechanisms, and how they may differ between and within TDPs, is warranted, enabling further optimization of the different TDP stimulation in clinical settings.

## Data availability

The datasets generated during and/or analyzed during the current study are available from the corresponding author on reasonable request.

## Acknowledgments

This work was supported by investigator-initiated grant from Boston Scientific Neuromodulation Corp. to CYS.

## Author contributions

M.M.E. conducted the study and contributed to the design of the study; C.Z., contributed to the design of the study, analyzed the data, and helped to draft the manuscript; K.J. analyzed the data and helped to draft the manuscript; V.R. contributed to data analysis; R.E. contributed to the design of the study and helped to draft the manuscript; C.Y.S. contributed to the conception and design of the study, interpretation of the data, and drafted the manuscript and revising it critically. The authors approved the final version of the manuscript.

## Competing interests

R.E. and C.Z. are employees of Boston Scientific Neuromodulation Corp. M.M.E., K.J., V.R., and C.Y.S. have no conflicts of interest to disclose.

## Notes

### Competing Interest Statement

The authors have declared no competing interest.

