## Supplemental Figure 1 for "Spinal Cord Stimulation using time-dynamic pulses achieves faster and longer reversal of allodynia compared to tonic pulses in a rat model of neuropathic pain"

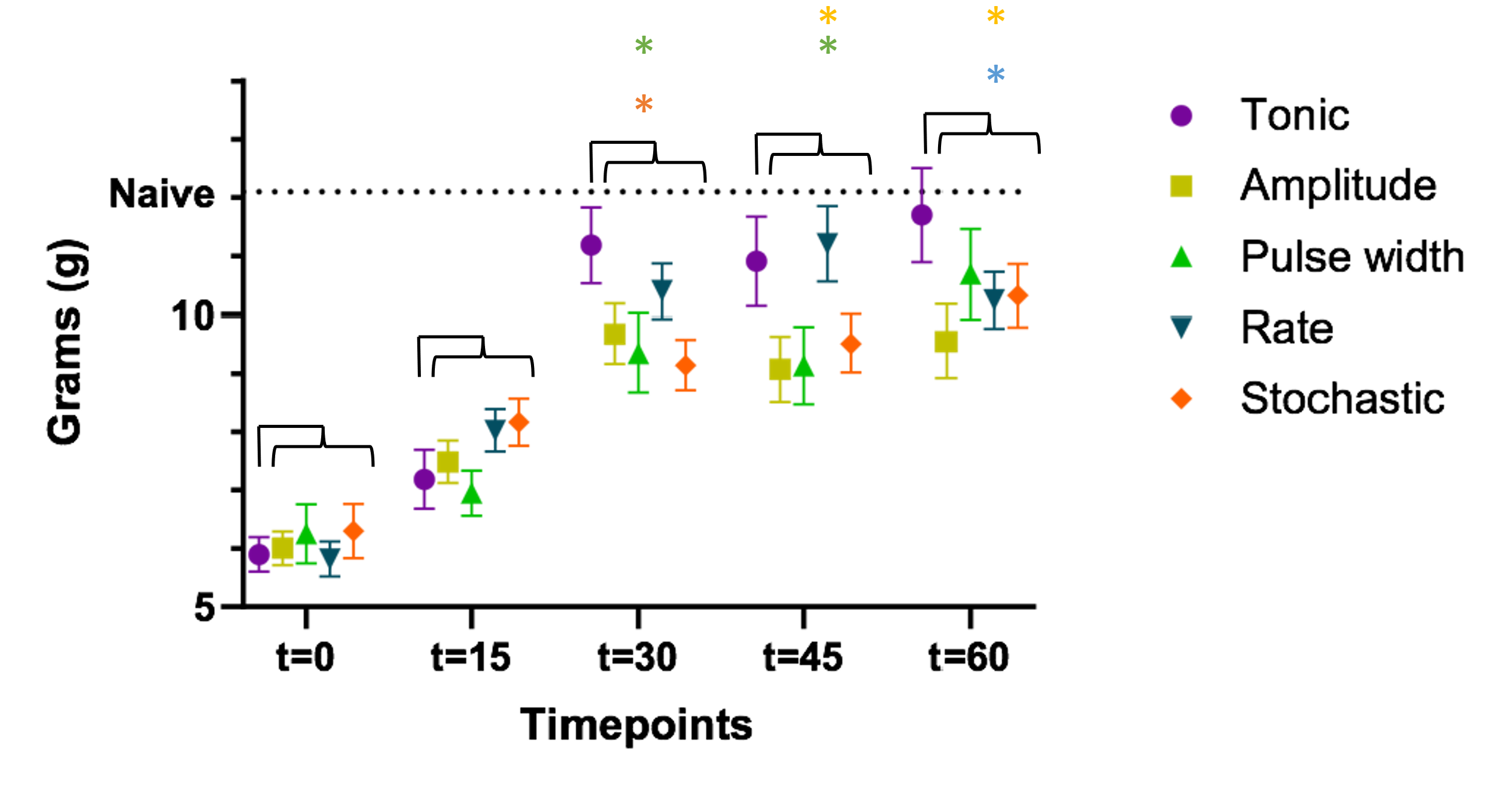
**Supplementary**

Supplementary Figure 1. Comparison of PWT time course across 5 different SCS patterns (n=23 rats). Tonic stimulation results in significantly higher PWT compared to amplitude and stochastic modulation at t=30, 45, and 60 min after SCS onset.
